# Biological management of fusarium wilt on tomato caused by *Fusarium oxysporum* f. sp. *lycospersici* by some plant growth-promoting bacteria

**DOI:** 10.1101/2020.08.21.262212

**Authors:** Halima Z. Hussein, Shaker I. Al-Dulaimi

## Abstract

Chemical approaches have been applied to combat Fusarium wilt disease for a long time. Even though pesticides are effective in controlling the disease, they continue to damage the environment. Environmental-friendly approaches to manage plant disease are the goal of many studies recently. This study was conducted to assess the efficacy of some bio-agents in induction of systemic resistance in tomato plants as a management approach of Fusarium wilt disease caused by *Fusarium oxysporum* f.sp. *lycopersici* (FOL) under condition Plastic house. Results of the plastic house experiments showed that all treatments in decreased Fusarium disease percentage and severity on tomato, two bacterial combinations (Streptomyces sp. (St) and *Pseudomonas fluorescence* (Pf)) decreased the infection percentage and disease severity with 16.6% and 8.3%, respectively. Treatment with St reduced the infection percentage and disease severity with 33.3% and 22.8%, while the Pf treatment showed 41.6% and 31.2% reduction in infection percentage and disease severity, compared to 100% and 91.6% in the control treatment. Results of induced systemic resistance (ISR) biochemical indicators showed significant differences in tomato plants. Peroxidase and Phenylalanine-Ammonia-Lyase (PAL) activity and the Phenol content increased significantly 14 days after treatments compared to the control treatment, which contains only the fungal pathogen FOL.

## Introduction

Tomato (*Solanum lycopersicum*) belongs to the Family Solanaceae, which is one of the world’s most widely cultivated vegetable crops (Srivastava *el at*, 2010; McGovern, 2015). Low yield of tomato is attributed to its susceptibility to several pathogenic fungi, bacteria, viruses and nematodes which are major constraints to tomato cultivation such as Fusarium wilt, gray mold, early blight, tomato leaf curl disease, bacterial wilt, damping off and Verticillium wilt (Al-Ani et al., 2011c). Among these, *Fusarium oxysporum* f.sp. *lycopersici* (FOL) (Sacc.) W.C.Synder and H.N. Hans, incident of vascular wilt of tomato, alone cause 30-40% yield loss and in India, under adverse weather conditions, the losses may reach as high as 80% (Bawa, 2016; Nirmaladevi *et al*, 2016; Sidharthan *et al*, 2018).

The pathogen invades the root epidermis and extends into the vascular tissue. It colonizes the xylem vessels producing mycelium and conidia. The characteristic wilt symptoms appear as a result of severe water stress, mainly due to vessel clogging (Kennelly, 2007; Girhepuje and Shinde, 2011; Ramanathan *et al*, 2010; McGovern, 2015). Three physiological races (1, 2, and 3) of the pathogen are distinguished by their specific pathogenicity to tomato cultivars (Reis and Boiteux, 2007; Biju *et al*, 2017). Since *F*.*oxysporum* f.sp. *lycopersici* (FOL) is an asexual fungus, genetic exchange occurs via somatic fusion and hetreokaryon formation between vegetative compatible strains (Leslie, 1993).

Chemical pesticides have been used to combat Fusarium wilt disease as well as polluting the environment and natural sources such as water, air and soil. In view of the negative effects of pesticides, researchers have sought to use alternative factors to reduce pathogens. Many biological agents have been proved as successful ecofriendly and viable alternative approaches to manage different plant diseases (Al-Ani et al., 2012, 2011a, 2011b, 2013; Al-Ani and Adhab 2012, 2013). These strategies aim to replace chemical pesticides with To be a good choice, Plant Growth Promotion Rhizobacteria (PGPR) was proposed for the management of plant diseases, PGPR is a useful microorganism found in an area around the roots that promotes growth and provides protection to plants such as nitrogen stabilization, phosphorus thawing, production of iron-bearing materials, volatile organic compounds, and reduction of disease effect such as the production of disintegration enzymes of fungal cell walls such as Chitinase, Glucanase, Protease, Lipase and Amylases which stimulates induced systemic resistance (ISR) (Vejan et al, 2016; Zouari et al, 2016; Mhlongo et al, 2018).

The objective of the study was to evaluate the effectiveness of PGPR bacteria in reducing the incidence of pathogenic fungi and to investigate their effectiveness in stimulating systemic resistance in the plant and increase plant defenses through the induction of Peroxidase and Phenylalanine-Ammonia-Lyase as well as the total Phenol content of the plant.

## Materials and methods

### Pathogenicity test

This experiment was conducted in the Glass House of the Department of Plant Protection / College of Agricultural Engineering Sciences / University of Baghdad. Isolate from the pathogen *Fusarium oxysporum* f.sp. *lycopersici* molecularly diagnosed, were obtained of the Mycotoxins Laboratory characterization morphological and restored molecular characterization using PCR technique, the sequence obtained was dispensed into GenBank under accession number MH458918 (Al-Dulaimi and Hussein, 2019).

Used sterile soil 2:1 for Pathogenicity test fungal FOL on seed tomato, were placed in pots capacity 1 kg and added half plates a 6-days-old fungal mycelia per pot then cover to 3 days and sowing 10 seed tomato surface sterile per pot and leave for 10 days. Germination percentage was calculated according to:

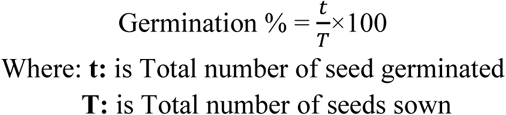

### Isolate PGPR bacteria and test their antifungal ability against FOL

Soil samples were collected from an area around the roots of tomato plants that were selected on the basis of good vegetative growth of plants from fields in the provinces of Baghdad and Anbar. Soil samples (1 g) were serially diluted at 1:10, in vitro antagonistic activity of PGPR against F.oxysporum f. sp. lycopersici. The experiment was replicated three times and the zone of inhibition was measured according to the following formula described in (Raspor *et al*, 2016).

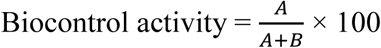

Where **A**: Is the area of inhibition, the distance between the biological control agent line and the end of the fungal growth, **B**: Fungal expansion towards the biological control agent line

### Morphological identification of PGPR

#### 1 Identification of *Streptomyces* spp

The purified Streptomyces spp. isolates were characterized to genus level according to Bergey’s Manual of Determinative Bacteriology (Breed *et al*, 1957).

#### 2 Identification *P. fluorescence*

The isolate was diagnosed at the Central Public Health Laboratory In the Iraqi Ministry of Health using the Analytical Profile Index 20 Enterobacteriaceae (API 20E).

### The efficacy of PGPR bacteria in resistance to pathogenic fungi *Fusarium oxysporum* f.sp. *lycopersici*

the experiment was carried out in the plastic house of the Department of Plant Protection /College of Agricultural Engineering Sciences / University of Baghdad, Sterilize mixed soil with steam sterilizer at 121°C and pressure 1.5 kg/cm^2^ for 20 minutes for Twice consecutive. It was then distributed in plastic pots diameter 15 cm (And at a rate of 1 kg/pot). 30-day tomato seedlings were planted a variety agro-TIP With four seedlings per pot The bacterial inoculum (PGPR) *Streptomyces* sp. (St) (10 ml 60×10^8^) and *P*.*fluorescens* (Pf) (10 ml 50×10^8^) was added on the same day of planting. The disease-causing the *F. oxysporum* f.sp. *lycopersici* strain was grown and maintained on a PDA plate and after 6-days of complete fungal growth, the fungus was cut in equal pieces, applied to root epithelial tissues and covered with soil, The control and treated plants were kept in dark condition at relatively high humidity level of 80% for 4-days in growth chamber in order to further exploit the pathogenic impact. The pots were arranged in a CRD in a set of five replicates. The following treatments were examined:

1. **Control**: (neither inoculated with *F*.*oxysporum* f.sp. *lycopersici* nor with microbial treatment).
2. **FOL**: (inoculated with *F*.*oxysporum* f.sp. *lycopersici*).
3. **St**: (inoculated with *Streptomyces* sp.).
4. **Pf**: (inoculated with *P*.*fluorescens*).
5. **FOL** + **St**
6. **FOL** + **Pf**
7. **St** + **Pf**
8. **FOL** + **St** + **Pf**

### Preparation of enzyme activities and Phenol content

The efficacy of Peroxidase enzyme as reported by Hammerschmidt *et al* (1989), The efficacy of the enzyme PAL according to the method mentioned by Dickerson *et al*(1984), Phenol content according to Rishi *et al* (2008). After 7, 14 and 21-days of pathogen inoculation, It has been effectively measured (ISR) in tomato plants is Peroxidase, PAL enzyme and Phenol content.

### Percentage of disease severities and infection

Symptom severity of the shoot system of the plants was assessed (6-weeks after pathogen inoculation) using the following index (Souza *et al*, 2010): **0:** no symptoms, **1:** yellowing 1 – 25% of the leaves near the stem base, **2:** Yellowing and wilt 26-50% of leaves with a simple brown discoloration in xylem vessels, **3:** Yellowing and wilt 51 – 75% of the leaves with dark brown coloring in xylem vessels, **4:** Wilt and die 76-100% of leaves.

The disease severity index was calculated using the formula (Mckinney, 1923).

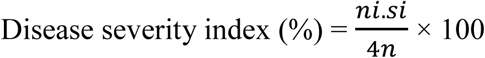

And Infection Percentage was calculated using the formula:

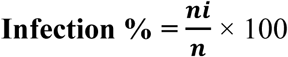

where ***ni***: is the number of plants affected by each degree of severity.

***si***: the degree of severity of the attack (0 – 4).

***n***: the total number of plants used for each energy level applied.

### Statistical analysis

The data were analyzed by ANOVA using SAS statistical program (SAS, 2012). Results were compared using the least significant difference (LSD) test at P 5%.

## Results

### Pathogenicity test

The result of the experiment showed that the isolate of pathogenic fungi resulted in a significant reduction in the percentage of germination of tomato seeds. The germination rate was 16.55% compared to the control treatment (without the presence of pathogenic fungus) with a germination rate of 100% (Table 1).

**Table 1.** Test the pathological potential of FOL on tomato seeds in the pots

| Treatment | Germination% |
| --- | --- |
| Control | 100 |
| Treatment with pathogen | 16.55 |
| L.S.D* | 24.49 |

• Each number represents the rate of three replicates

Due to the ability of the pathogen FOL to cause high infection rate and prevent germination of seeds significantly in the experience of the pots to the ability to secrete the enzymes that analyze the walls of cells such as Chitinase and Cellulase, which break down chitin and cellulose as well as have an important role in the virulence of the causative agent, And its production of toxic metabolic materials such as Dehydro Fusaric acid and Lycomarsmin and Fusaric acid, which affect the permeability of membranes of infected plant cells and reduce respiratory rates in the plant cell and cause imbalance in other plant biological processes (Burgess *et al*, 1998; Langcalces and Drysdale, 2005; Calero-Nieto *et al*, 2007; Ramanathan *et al*, 2010).

### Test of Antifungal activities of PGPR

45-isolates of PGPR were encountered from soil fields in Baghdad and Anbar planted with tomato, Dual culture test showed that two isolates of soil PGPR revealed the highest antifungal activity against *F*.*oxysporum* f.sp. *lycopersici* as 100 % inhibition of colony growth as follows: St, Pf.

### Identification of the effective soil PGPR

#### 1 Identification of Streptomyces spp.

Morphological and chemotaxonomy identification: The results of the microscopic examination of bacterial isolation of St on PDA, ISP2 and ISP4 showed that the colonies were white on the PDA medium, and red to pink on ISP2 and ISP4 medium has a dusty surface structure and be mycelium like hyphae fungus, a characteristic of bacteria *Streptomyces* spp. as well as able to produce spores and these spores are responsible for red color (Kieser *et al*, 2000; Kampfer, 2006).

#### 2 Identification *P. fluorescence*

The results of the diagnosis of F11 isolation were shown using technology Analytical Profile Index 20 Enterobacteriaceae (API 20E), by comparing the numbers obtained are (2002004) with the numbers mentioned in the manufacturer’s manual, bacteria can identify the gene and species of bacteria to be diagnosed. It has been shown to belong to *Pseudomonas fluorescence* an important species of the PGPR.

### Evaluation of the efficacy of PGPR bacteria in control of Fusarium wilt disease in tomato plant

All transactions (Table 2) achieved a significant decrease in the percentage of disease severities and infection of Fusarium wilt disease in a tomato plant and in varying degrees, As shown by treatment between the isolates of the bacteria with the presence of pathogen (FOL + St + Pf) has best rate of prevention of infection, as the percentage of disease severities and infection 16.6% and 8.3%. Followed by the treatment of bacterial isolation Streptomyces sp. with pathogenic fungi (FOL + St) scored 33.3% and 22.8% and treatment (FOL + Pf) it reached 41.6% and 31.2%, compared with the treatment of pathogenic fungi alone (FOL) scored 100 and 91.6%, respectively.

**Table 2.** Efficacy of PGPR bacteria in control of Fusarium wilt disease

| treatments | Infection % | Disease severities % |
| --- | --- | --- |
| Control | 0 | 0 |
| FOL | 100 | 91.6 |
| FOL + St | 33.3 | 22.8 |
| FOL + Pf | 41.6 | 31.2 |
| FOL + St + Pf | 16.6 | 8.3 |
| L.S.D | 3.4 | 2.2 |

• Each number represents the rate of three replicates

The decrease in the percentage of disease severities and infection of Fusarium wilt disease by the efficiency of the agents used which has two directions in resisting pathogens. The first trend is the direct effect on the pathogen it inhibits the growth of mycelium radial and inhibits the germination of the spores and the ability of bacteria to produce antibiotics and disintegration enzymes such as Protease, Chitinase, Amylase, and Lipase, and produce HCN.

### Efficacy of PGPR bacteria to induce systemic resistance in tomato

There was an increased activity of defense-related enzymes Peroxidase, PAL and Phenol content due to the application of bacteria PGPR when challenge inoculated with *F*.*oxysporum* f.sp. *lycopersici* (Fig 1,2 and 3). Treatments used showed high efficacy to affect Peroxidase, PAL and Phenol content when biochemical indicators activity 14 after days of the pathogen added. treatment bacteria *Streptomyces* sp. With pathogenic fungus (FOL + St + Pf) which scored 63.8, 32.6 and 3.8. Whereas treatment *Streptomyces* sp. with pathogenic fungus (FOL + St) which scored 48.6, 27.4 and 3.2, then treatment *P*.*fluorescens* With pathogenic fungus (FOL+ Pf) scored 43.6, 26.3 and 3.1. Compared to pathogenic fungus alone control which scored 22.3, 16.6 and 2.0 enzyme activity, respectively.

**Fig 1.**
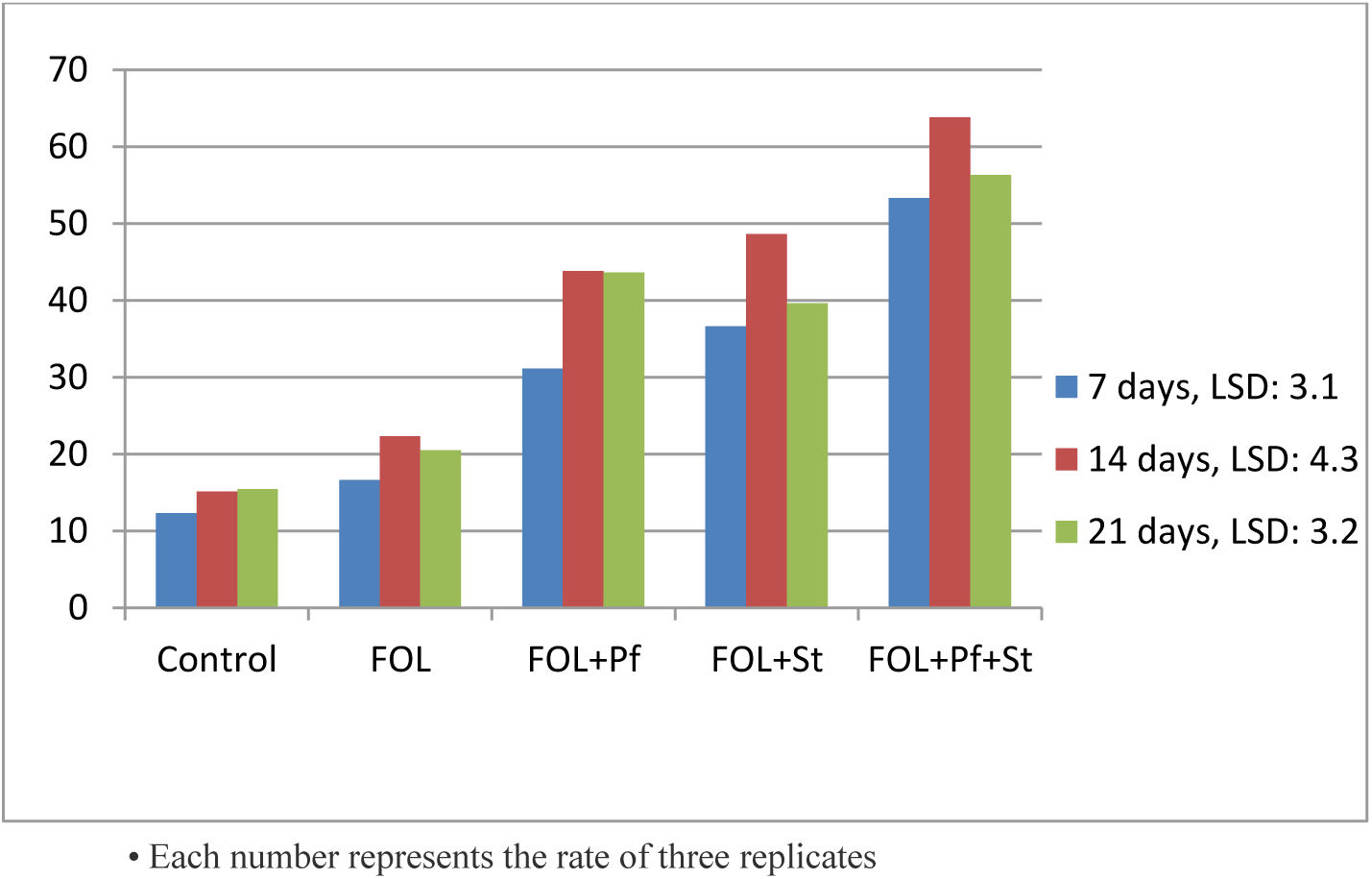
Peroxidase activity

**Fig 2.**
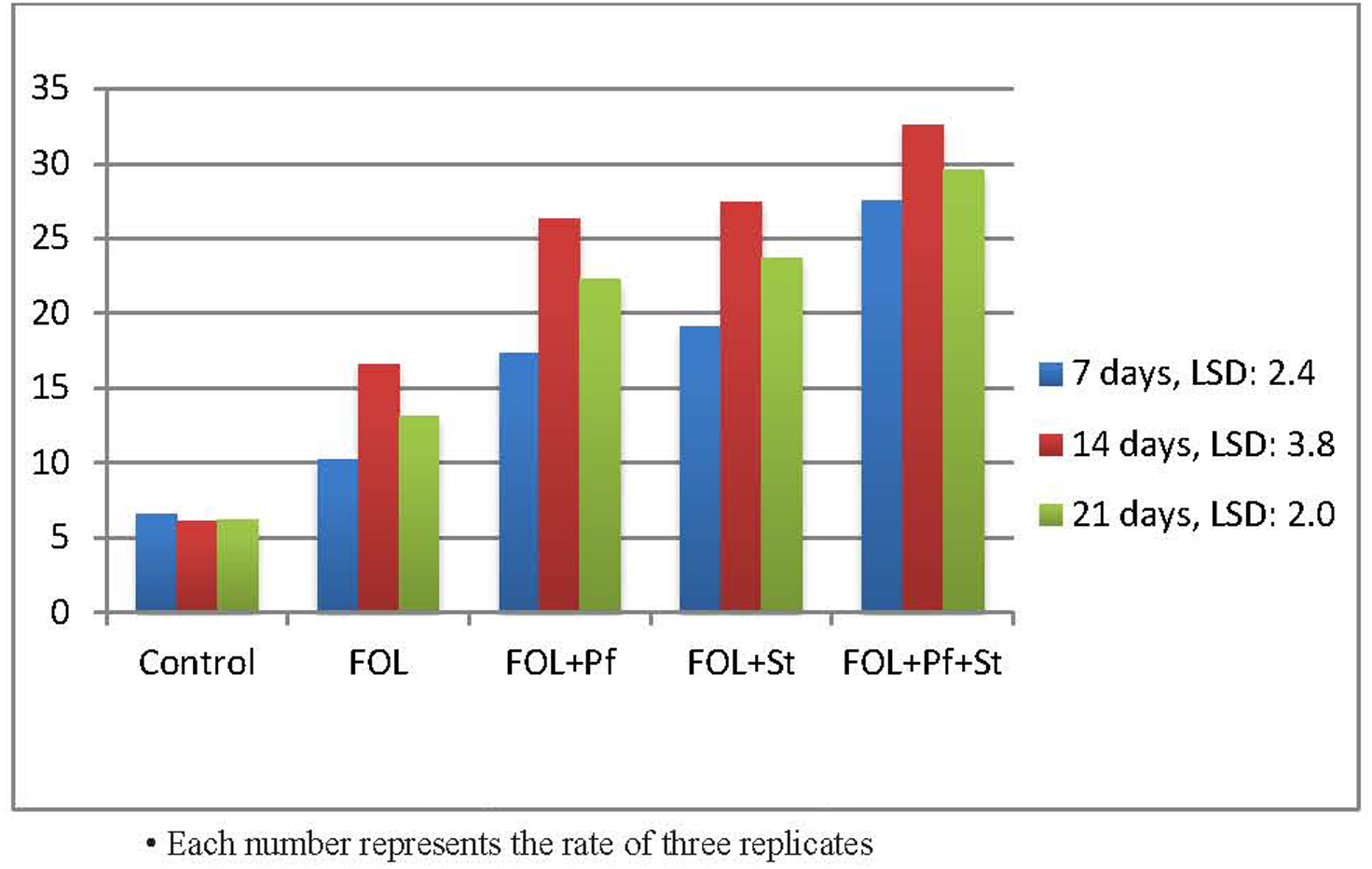
PAL activity

**Fig 3.**
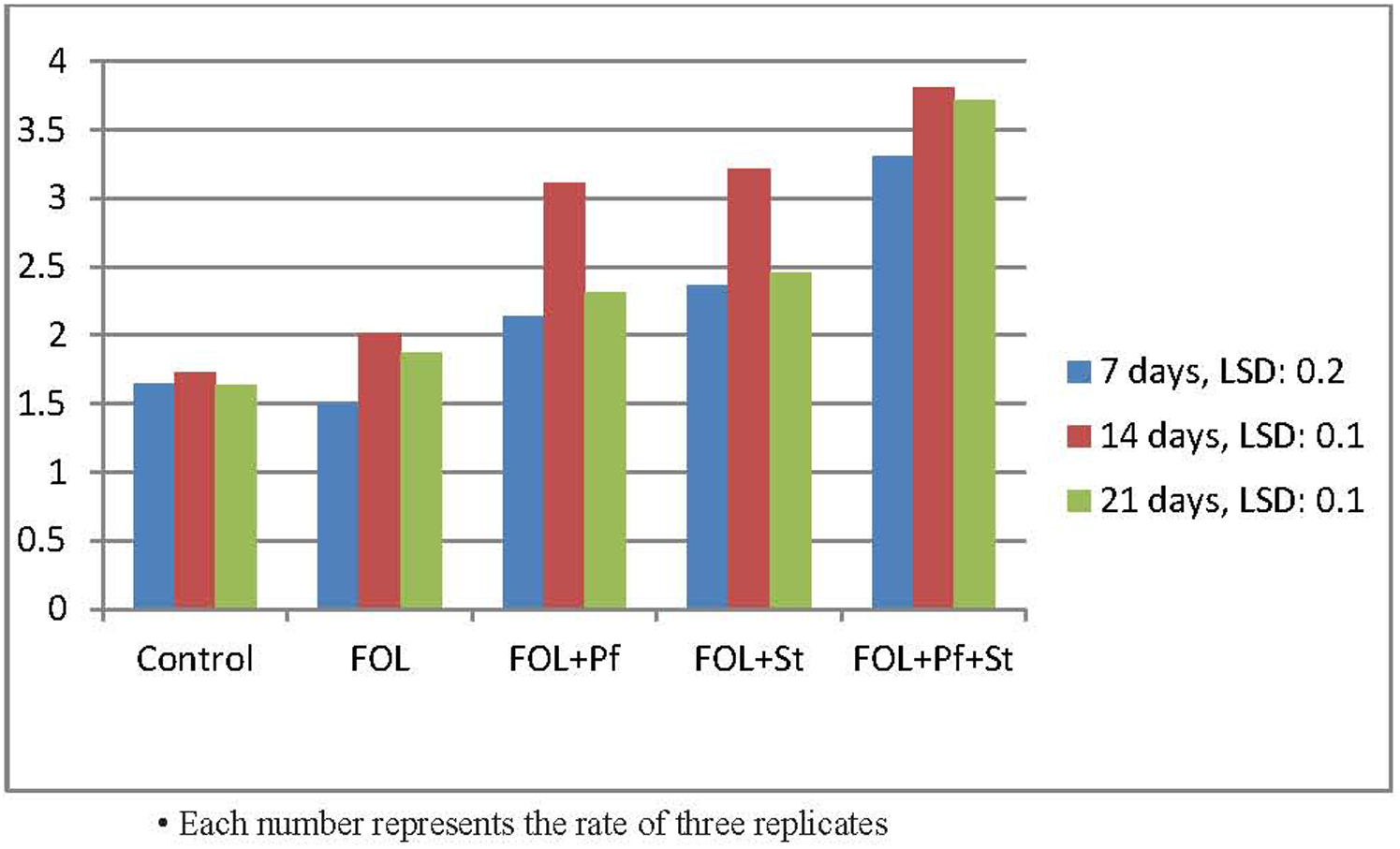
Phenol content

Peroxidase and PAL enzyme is stimulated in plant tissues when exposed to pathogens or inducers and has a crucial role in determining the level of host resistance. It is a major enzyme in the biosynthesis of lignin, the deposition of the Suberin, and the intensity of the cell wall which increased structural defenses (Thakker *et al*, 2013; Cass *et al*, 2015; Chen *et al*, 2017).

